# Conventional and hyperspectral time-series imaging of maize lines widely used in field trials

**DOI:** 10.1101/169045

**Authors:** Zhikai Liang, Piyush Pandey, Vincent Stoerger, Yuhang Xu, Yumou Qiu, Yufeng Ge, James C. Schnable

## Abstract

Maize (*Zea mays* ssp. *mays*) is one of three crops, along with rice and wheat, responsible for more than 1/2 of all calories consumed around the world. Increasing the yield and stress tolerance of these crops is essential to meet the growing need for food. The cost and speed of plant phenotyping is currently the largest constraint on plant breeding efforts. Datasets linking new types of high throughput phenotyping data collected from plants to the performance of the same genotypes under agronomic conditions across a wide range of environments are essential for developing new statistical approaches and computer vision based tools. A set of maize inbreds – primarily recently off patent lines – were phenotyped using a high throughput platform at University of Nebraska-Lincoln. These lines have been previously subjected to high density genotyping, and scored for a core set of 13 phenotypes in field trials across 13 North American states in two years by the Genomes to Fields consortium. A total of 485 GB of image data including RGB, hyperspectral, fluorescence and thermal infrared photos has been released. Correlations between image-based measurements and manual measurements demonstrated the feasibility of quantifying variation in plant architecture using image data. However, naive approaches to measuring traits such as biomass can introduce nonrandom measurement errors confounded with genotype variation. Analysis of hyperspectral image data demonstrated unique signatures from stem tissue. Integrating heritable phenotypes from high-throughput phenotyping data with field data from different environments can reveal previously unknown factors influencing yield plasticity.

## Data Description

### Background

The green revolution created a significant increase in the yields of several major crops in the 1960s and 1970s, dramatically reducing the prevalence of hunger and famine around the world, even as population growth continued. One of the major components of the green revolution was new varieties of major grain crops produced through conventional phenotypic selection with higher yield potentially. Since the green revolution, the need for food has continued to increase, and a great deal of effort in the public and private sectors is devoted to developing crop varieties with higher yield potential. However, as the low hanging fruit for increased yield vanish, each new increase in yield requires more time and resources. Recent studies have demonstrated that yield increases may have slowed or stopped for some major grain crops in large regions of the world^1^. New approaches to plant breeding must be developed if crop production continues to grow to meet the needs of an increasing population around the world.

The major bottleneck in modern plant breeding is phenotyping. Phenotyping can be used in two ways. Firstly, by phenotyping a large set of lines, a plant breeder can identify those lines with the highest yield potential and/or greatest stress tolerance in a given environment. Secondly, sufficiently detailed phenotyping measurements from enough different plants can be combined with genotypic data to identify regions of the genome of a particular plant species which carry beneficial or deleterious alleles. The breeder can then develop new crop varieties which incorporate as many beneficial alleles and exclude as many deleterious alleles as possible. Phenotyping tends to be expensive and low throughput, yet as breeders seek to identify larger numbers of alleles each with individually smaller effects, the amount of phenotyping required to achieve a given increase in yield potential is growing. High throughput computer vision based approaches to plant phenotyping have the potential to ameliorate this bottleneck. These tools can be used to precisely quantify even subtle traits in plants and will tend to decrease in unit cost with scale, while conventional phenotyping, which remains a human labor intensive processes, does not.

Several recent pilot studies have applied a range of image-processing techniques to extract phenotypic measurements from crop plants. RGB (R: Red channel; G: Green channel; B: Blue channel) camera technology, widely used in the consumer sector, has also been the most widely used tool in these initial efforts at computer vision based plant phenotyping^2–5^. Other types ofcameras including fluoresence^6,7^ and NIR (near-infrared)^6,8,9^ have also been employed in high throughput plant phenotyping efforts, primarily in studies of the response of plant to different abiotic stresses.

However, the utility of current studies is limited in two ways. Firstly, current analysis tools can extract only a small number of different phenotypic measurements from images of crop plants. Approximately 150 tools for analyzing plant image data are listed in a field specific database, however the majority of these are either developed specifically for *Arabidopsis thaliana* which is a model plant, or are designed specifically to analyze images of roots^10^. Secondly, a great deal of image data is generated in controlled environments, however, there are comparatively few attempts to link phenotypic measurements in the greenhouse to performance in the field. However, one recent report in maize suggested that more than 50% of the total variation in yield under field conditions could be predicted using traits measured under controlled environments^5^.

Advances in computational tools for extracting phenotypic measurements of plants from image data and statistical models for predicting yield under different field conditions from such measurements requires suitable training datasets. Here, we generate and validate such a dataset consisting of high throughput phenotyping data from 32 distinct maize (*Zea mays*) accessions drawn primarily from recently off-patent lines developed by major plant breeding companies. These accessions were selected specifically because paired data from the same lines exists for a wide range of plant phenotypes collected in 54 distinct field trials at locations spanning 13 North American states or provinces over two years^11^. This extremely broad set of field sites captures much of the environmental variation among areas in which maize are cultivated with total rainfall during the growing season ranging from 133.604 mm to 960.628 mm (excluding sites with supplemental irrigation) and peak temperatures during the growing season ranging from 23.°C to 34.9°C. In addition, the same lines have been genotyped for approximately 200,000 SNP markers using GBS^11^. Towards these existing data, we added RGB, thermal infra-red, fluorescent and hyperspectral images collected once per day per plant, as well as detailed water-use information (single day, single plant resolution). At the end of the experiment, 12 different types of ground-truth phenotypes were measured for individual plants including destructive measurements. A second experiment focused on interactions between genotype and environmental stress, collecting the same types of data described above from two maize genotypes under well watered and water stressed conditions^12^. We are releasing this curated dataset of high throughput plant phenotyping images from accessions where data on both genotypic variation and and agronomic performance under field conditions is already available. All data was generated using a Lemnatec designed high throughput greenhouse-based phenotyping system constructed at the University of Nebraska-Lincoln. This system is distinguished from existing public sector phenotyping systems in North America by both the ability to grow plants to a height of 2.5 meters and the incorporation of a hyperspectral camera^9^. Given the unique properties described above, this comprehensive data set should lower the barriers to the development of new computer vision approaches or statistical methodologies by independent researchers who do not have the funding or infrastructure to generate the wide range of different types of data needed.

## Methods

### 0.0.1 Greenhouse Management

All imaged plants were grown in the greenhouse facility of the University of Nebraska-Lincoln’s Greenhouse Innovation Center (Latitude: 40.83, Longitude: −96.69) between October 2nd, 2015 to November 10th, 2015. Kernels were sown in 1.5 gallon pots with Fafard germination mix supplemented with 1 cup (236 mL) of Osmocote plus 15-9-12 and one tablespoon (15 mL) of Micromax Micronutrients per 2.8 cubic feet (80 L) of soil. The target photoperiod was 14:10 with supplementary light provided by LED growth lamps from 07:00 to 21:00 each day. The target temperature of the growth facility was between 24 *–* 26°C. Pots were weighed once per day and watered back to a target weight of 5,400 grams from 10-09-2015 to 11-07-2015 and a target weight of 5,500 grams from 11-08-2015 to the termination of the experiment.

### 0.0.2 Experimental Design

A total of 156 plants, representing the 32 genotypes listed in Table 1 were grown and imaged, as well as 4 pots with soil but no plant which serve as controls for the amount of water lost from soil as a result of non-transpiration mechanisms (e.g. evaporation). The 156 plants plus control pots were arranged in a ten row by sixteen column grid, with 0.235 meter spacing between plants in the same row and 1.5 meters spacing between rows (Table 2). Sequential pairs of two rows were consisted of a complete replicate with either 31 genotypes and one empty control pot, or 32 genotypes. Within each pair of rows, genotypes were blocked in groups of eight (one half row), with order randomized within blocks between replicates in order to maximize statistical power to analyze within-greenhouse variation.

**Table 1.** 32 genotypes in maize phenotype map

| Genotype ID | Genotype | Source | Released Year |
| --- | --- | --- | --- |
| ZL1 | 740 | Novartis Seeds | 1998 |
| ZL2 | 2369 | Cargill | 1989 |
| ZL3 | A619 | Public Sector | 1992 |
| ZL4 | A632 | Public Sector | 1992 |
| ZL5 | A634 | Public Sector | 1992 |
| ZL6 | B14 | Public Sector | 1968 |
| ZL7 | B37 | Public Sector | 1971 |
| ZL8 | B73 | Public Sector | 1972 |
| ZL9 | C103 | Public Sector | 1991 |
| ZL10 | CM105 | Public Sector | 1992 |
| ZL11 | LH123HT | Holden's Foundation | 1984 |
| ZL12 | LH145 | Holden's Foundation | 1983 |
| ZL13 | LH162 | Holden's Foundation | 1990 |
| ZL14 | LH195 | Holden's Foundation | 1989 |
| ZL15 | LH198 | Holden's Foundation | 1991 |
| ZL16 | LH74 | Holden's Foundation | 1983 |
| ZL17 | LH82 | Holden's Foundation | 1985 |
| ZL18 | Mo17 | Public Sector | 1964 |
| ZL19* | DKPB80 | DEKALB Genetics | ? |
| ZL20 | PH207 | Pioneer Hi-Bred | 1983 |
| ZL21 | PHB47 | Pioneer Hi-Bred | 1983 |
| ZL22** | PHG35 | Pioneer Hi-Bred | 1983 |
| ZL23 | PHG39 | Pioneer Hi-Bred | 1983 |
| ZL24 | PHG47 | Pioneer Hi-Bred | 1986 |
| ZL25 | PHG83 | Pioneer Hi-Bred | 1985 |
| ZL26 | PHJ40 | Pioneer Hi-Bred | 1986 |
| ZL27 | PHN82 | Pioneer Hi-Bred | 1989 |
| ZL28 | PHV63 | Pioneer Hi-Bred | 1988 |
| ZL29 | PHW52 | Pioneer Hi-Bred | 1988 |
| ZL30 | PHZ51 | Pioneer Hi-Bred | 1986 |
| ZL31 | W117HT | Public Sector | 1982 |
| ZL32 | Wf9 | Public Sector | 1991 |

\* Not currently available for order. \*\* Genotype repre-
sented by only a single plant in the dataset.

**Table 2.**
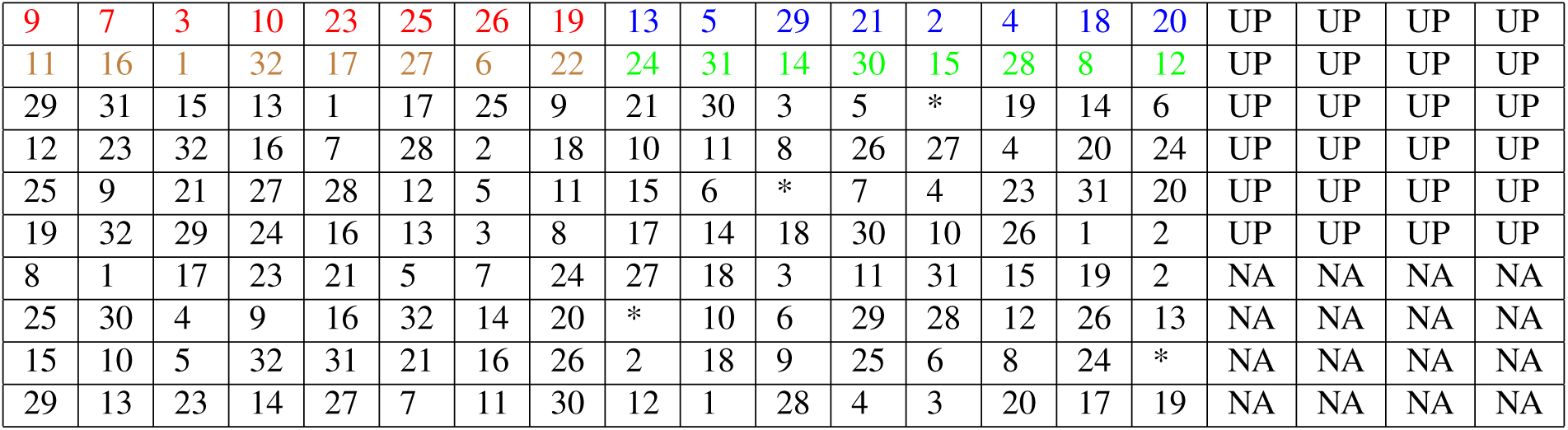
Experimental layout (ID: ZL1-ZL32). At the time this experiment was conducted, the total size of the UNL greenhouse system was ten rows by twenty columns. Positions marked with UP indicate pots filled with plants from an unrelated experiment, while positions marked with NA indicate pots which had no plants. The first complete replicate is shown in color, and the four incomplete blocks within the first replicate are marked in different colors. * marks empty pots within the experimental design.

|  |  |  |  |  |  |  |  |  |  |  |  |  |  |  |  |  |  |  |  |
| --- | --- | --- | --- | --- | --- | --- | --- | --- | --- | --- | --- | --- | --- | --- | --- | --- | --- | --- | --- |
| 9 | 7 | 3 | 10 | 23 | 25 | 26 | 19 | 13 | 5 | 29 | 21 | 2 | 4 | 18 | 20 | UP | UP | UP | UP |
| 11 | 16 | 1 | 32 | 17 | 27 | 6 | 22 | 24 | 31 | 14 | 30 | 15 | 28 | 8 | 12 | UP | UP | UP | UP |
| 29 | 31 | 15 | 13 | 1 | 17 | 25 | 9 | 21 | 30 | 3 | 5 | * | 19 | 14 | 6 | UP | UP | UP | UP |
| 12 | 23 | 32 | 16 | 7 | 28 | 2 | 18 | 10 | 11 | 8 | 26 | 27 | 4 | 20 | 24 | UP | UP | UP | UP |
| 25 | 9 | 21 | 27 | 28 | 12 | 5 | 11 | 15 | 6 | * | 7 | 4 | 23 | 31 | 20 | UP | UP | UP | UP |
| 19 | 32 | 29 | 24 | 16 | 13 | 3 | 8 | 17 | 14 | 18 | 30 | 10 | 26 | 1 | 2 | UP | UP | UP | UP |
| 8 | 1 | 17 | 23 | 21 | 5 | 7 | 24 | 27 | 18 | 3 | 11 | 31 | 15 | 19 | 2 | NA | NA | NA | NA |
| 25 | 30 | 4 | 9 | 16 | 32 | 14 | 20 | * | 10 | 6 | 29 | 28 | 12 | 26 | 13 | NA | NA | NA | NA |
| 15 | 10 | 5 | 32 | 31 | 21 | 16 | 26 | 2 | 18 | 9 | 25 | 6 | 8 | 24 | * | NA | NA | NA | NA |
| 29 | 13 | 23 | 14 | 27 | 7 | 11 | 30 | 12 | 1 | 28 | 4 | 3 | 20 | 17 | 19 | NA | NA | NA | NA |

### Plant imaging

The plants were imaged daily using four different cameras in separate imaging chambers. The four types of cameras were thermal infrared, fluorescence, conventional RGB, and hyperspectral^12^. Images were collected in the order that the camera types are listed in the previous sentence. On each day, plants were imaged sequentially by row, starting with row 1 column 1 and concluding with row 10, column 16 (Table 2).

Plants were imaged from the side at two angles offset 90 degrees from each other as well as a top down view. On the first day of imaging or when plants reached the two leaf stage of development, the pot was rotated so that the major axis of leaf phylotaxy was parallel to the camera in the PA0 orientation and perpendicular to the camera in the PA90 orientation. This orientation is consistent for all cameras and was not adjusted again for the remainder of the experiment. The fluorescence camera captured images with a resolution of 1038 *×* 1390 pixels and measures emission intensity at wavelengths between 500-750 nm based on excitation with light at 400-500 nm. Plants were imaged using the same three perspectives employed for the thermal infrared camera. The RGB camera captured images with a resolution of 2454 *×* 2056 pixels. Initially the zoom of the RGB camera in side views was set such that each pixel corresponds to 0.746 mm at the distance of the pot from the camera. Between 2015-11-05 and 2015-11-10, the zoom level of the RGB camera was reduced to keep the entire plant in the frame of the image. As a result of a system error, this same decreased zoom level was also applied to all RGB images taken on 2015-10-20. At this reduced zoom level, each pixel corresponds to 1.507 mm at the distance of the pot from the camera, an approximate 2x change. Plants were also imaged using the same three perspectives employed for the thermal infrared camera.

The hyperspectral camera captured images with a resolution of 320 horizontal pixels. As a result of the scanning technology employed, vertical resolution ranged from 494 to 499 pixels. Hyperspectral imaging was conducted using illumination from halogen bulbs (Manufacturer Sylvania, model # ES50 HM UK 240V 35W 25 * GU10). A total of 243 separate intensity values were captured for each pixel spanning a range of light wavelengths between 546nm-1700nm. Data from each wavelength was stored as a separate grayscale image.

### Ground Truth Measurement

Ground truth measurements were collected at the termination of data collection on November 11-12, 2015. Manually collected phenotypes included plant height, total number of visible leaves, number of total fully extended leaves, stem diameter at the base of the plant, stem diameter at the collar of the top fully extended leaf, length and width of top fully extended leaf, and presence/absence visible anthocyanin production in the stem. After these measurements, total above-ground fresh weight biomass was measured for four out of five replicates, resulting in the destruction of the plants. Ground truth data for the drought stressed subset of this dataset was collected following the procedure previously described in^12^.

### RGB image processing

Pixels covering portions of the plant were segmented out of RGB images using a green index ((2*×*G)/(R+B)). Pixels with an index value greater than 1.15^12^ were considered to be plant pixels. This method produced some false positive plant pixels within the reflective metal columns at the edge of the image. To reduce the impact of false positives, these areas were excluded from the analysis. Therefore, when plant leaves cross the reflective metal frame, some true plant pixels were excluded. If no plant pixels were identified in the image – often the case in the first several days when the plant had either not germinated or had not risen above the edge of the pot – the value was recorded as “NA” in the output file.

### Heritability analysis

A linear regression model was used to analyze the genotype effect (excluding genotype ZL22 which lacked replication) and greenhouse position effect on plant traits. The responses were modeled independently for each day as

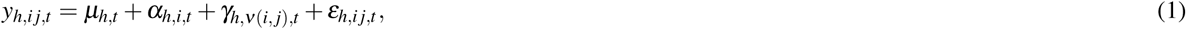

where the subscript *h* = 1,*…,*6 denotes the three responses extracted from the images: plant height, width and size for the two views 0 and 90 degree. The subscripts *i*, *j* and *t* denote the *j*th plant in the *i*th row and day *t*, respectively, and *n*(*i, j*) stands for the genotype at this pot. The parameters *α* and *γ* denote row effect and genotype effect, respectively. The error term is *ε*_h,i_*j,t*. Let SS_α,t_, SS_γ,t_ and SS_γ,t_ be the sum of squares of the regression model (1) for the row effect, genotype effect and the error at time *t*, respectively. Let SS_t_ = SS_α,t_ + SS_γ,t_ + SS_ε,t_ be the total sum of squares at time *t*. The heritability HR_t_ (2) of a given trait within this population was defined as the ratio of the genotype sum of squares over the sum of genotype and error sum of squares. For the estimate of the heritability of measurement error, the row effect term was replaced by a replicate effect (eachreplicate consisted of two sequential rows) with exclusion of ZL22 as only one plant of this genotype was grown.

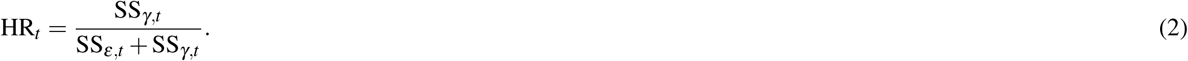

As the heritability index may change over the growth of the plant, an nonparametric smoothing method was provided for analyzing the time varying heritability of plants. The definition in (3) excludes the variation brought by the greenhouse row effect, which can be considered as the percentage of the variation in plant response that can be explained by the genotype effect after adjusting the environmental effect. To compare with this definition of heritability (2), the response in the model without considering the row effect was constructed as

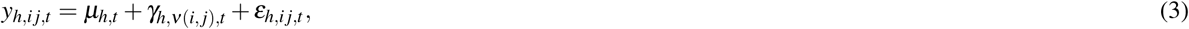

where similarly as (1), *n*(*i, j*) is the genotype of the *j*th plant in the *i*th row. Let 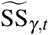 and 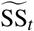 total sum of squares under (4). The classical heritability is defined as

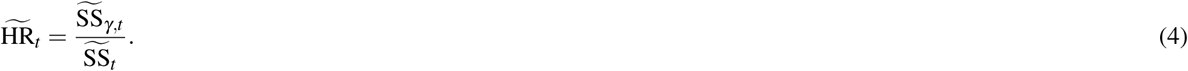

### Hyperspectral image processing

Two methods and thresholds were used to extract plant regions of interest from hyperspectral images. First, the commonly used NDVI (normalized difference vegetation index) formula was applied to all pixels using the formula (*R*_750nm-_*R*_705nm_)/(*R*_750nm_+*R*_705nm_), and pixels with a value greater than 0.25 were classified as originating from the plant^13^. Second, based on the difference in reflectance between stem and leaves at wavelengths of 1056nm and 1151nm, the stem was segmented from other part of plants by selecting pixels where (*R*_1056nm_/*R*_1151nm_) produced a value greater than 1.2. Leaf pixels were defined as pixels identified as plant pixels based on NDVI but not classified as stem pixels. In addition to the biological variation between individual plants, overall intensity variation existed both between different plants imaged on the same day and the same plant on different days as a result of changes in the performance of the lighting used in the hyperspectral imaging chamber. To calibrate each individual image and make the results comparable, a python script (hosted on Github; see code availability section) was used to normalize the intensity values of each plant pixel using data from the non-plant pixels in the same image.

In order to visualize variation across 243 separate wavelength measurements across multiple plant images, we used a PCA (Principal Component Analysis) based approach. After the normalization described above, PCA analysis of intensity values for individual pixels was conducted. PCA values of each individual plant pixel per analyzed plant were translated to intensity values using the formula [x-min(x)]/[max(x)-min(x)]. False color RGB images were constructed with the values for the first principal component stored in the red channel, the second principal component in the green channel and the third principal component stored in the blue channel.

### Fluorescence image processing

A consistent area of interest was defined for each zoom level to exclude the pot and non-uniform areas of the imaging chamber backdrop. Within that area, pixels with an intensity value greater than 70 in the red channel were considered to be plant pixels. The aggregate fluorescence intensity was defined as the sum of the red channel intensity values for all pixels classified as plant pixels within the region of interest, and the mean fluorescence intensity as the aggregate fluorescence intensity value divided by the number of plant pixels within the region of interest.

### Plant biomass prediction

Two methods were used to predict plant biomass. The first was a single variable model based on the number of zoom 3level adjusted plant pixels identified in the two RGB side view images on a given day. The second was a multivariate model based upon the sum of plant pixels identified in the two RGB side views, sum of plant pixels identified in the two RGB side views plus the RGB top view, aggregate fluorescence intensity in the two side views, aggregate fluorescence intensity in the two side views plus the top view, number of plant stem pixels identified in the hyperspectral image and number of plant leaf pixels identified in the hyperspectral image. Traits were selected to overlap with those employed by (Chen et al^14^) where possible. This multivariate dataset was used to predict plant biomass using linear modeling as well as MARS, Random Forest and SVM^14^.MARS analysis was performed using the R package earth^15^, Random Forest with the R package randomForest^16^ and SVM with the R package e1071^17^.

### Data Validation and quality control

### Validation against ground truth measurements

A total of approximately 500 GB of image data was initially generated by the system during the course of this experiment consisting of RGB images (51.1%), fluorescence images (4.3%), and hyperspectral images (44.6%). A subset of the RGB images within this dataset were previously analyzed in^18^, and were made available for download from http://plantvision.unl.edu/dataset under the terms of the Toronto Agreement. To validate the dataset and ensure plants had been properly tracked through both the automated imaging system and ground truth measurements, a simple script was written to segment images into plant and not-plant pixels (Figure 1). Source codes for all validation analysis are posted online (https://github.com/shanwai1234/Maize Phenotype Map).

**Figure 1.**
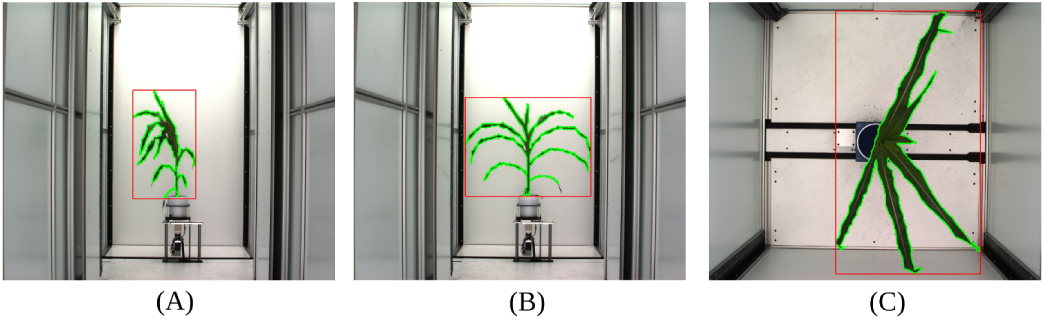
Segmentation of images into plant and not plant pixels for one representative plant (Path to this image in the released dataset: Genotype ZL019 *- >* Plant 008-19 *- >* Image Type *- >* Day 32). The area enclosed by green border is composed of pixels scored as “plant", the area outside the green border s composed of pixels scored as “not-plant". Minimum bounding rectangle of plant pixels is shown in red. (A) Side view, angle 1; (B) Side view, 90 degree rotation relative to A; (C) Top View.

Based on the segmentation of the image into plant and non-plant pixels, plant height was scored as the y axis dimension of the minimum bounding box. Plant area was scored as the total number of plant pixels observed in both side view images after correcting for the area of each pixel at each zoom employed (See Methods). Similar approaches to estimate plant biomass have been widely employed across a range of grain crop species including rice^19^, wheat^20^, barley^20,21^, maize^12^, sorghum^22^ and seteria^9^. Calculated values were compared to manual measurements of plant height and plant fresh biomass which were quantified using destructive methods on the last day of the experiment. In both cases manual measurements and image derived estimates were highly correlated, although the correlation between manual and estimated height was greater than the correlation between manually measured and estimated biomass (Figure 2A,B). Using the PlantCV software package^23^,equivalent correlations between estimated and ground truth biomass were obtained (r=0.91). Estimates of biomass using both software packages were more correlated with each other (r=0.96) than either was with ground truth measurements. This suggests that a significant fraction of the remaining error is the result of the expected imperfect correlation between plant size and plant mass, rather than inaccuracies in easimating plant size using individual software packages. Recent reports have suggested that estimates of biomass incorporating multiple traits extracted from image data can increase accuracy^14^.We tested the accuracy of biomass prediction of four multivariate estimation techniques on this dataset (see Methods). The correlation coefficient (r value) of the estimated biomass measures with ground truth data was 0.949, 0.958, 0.925 and 0.951 for multivariate linear model, MARS, Random Forest and SVM respectively.

**Figure 2.**
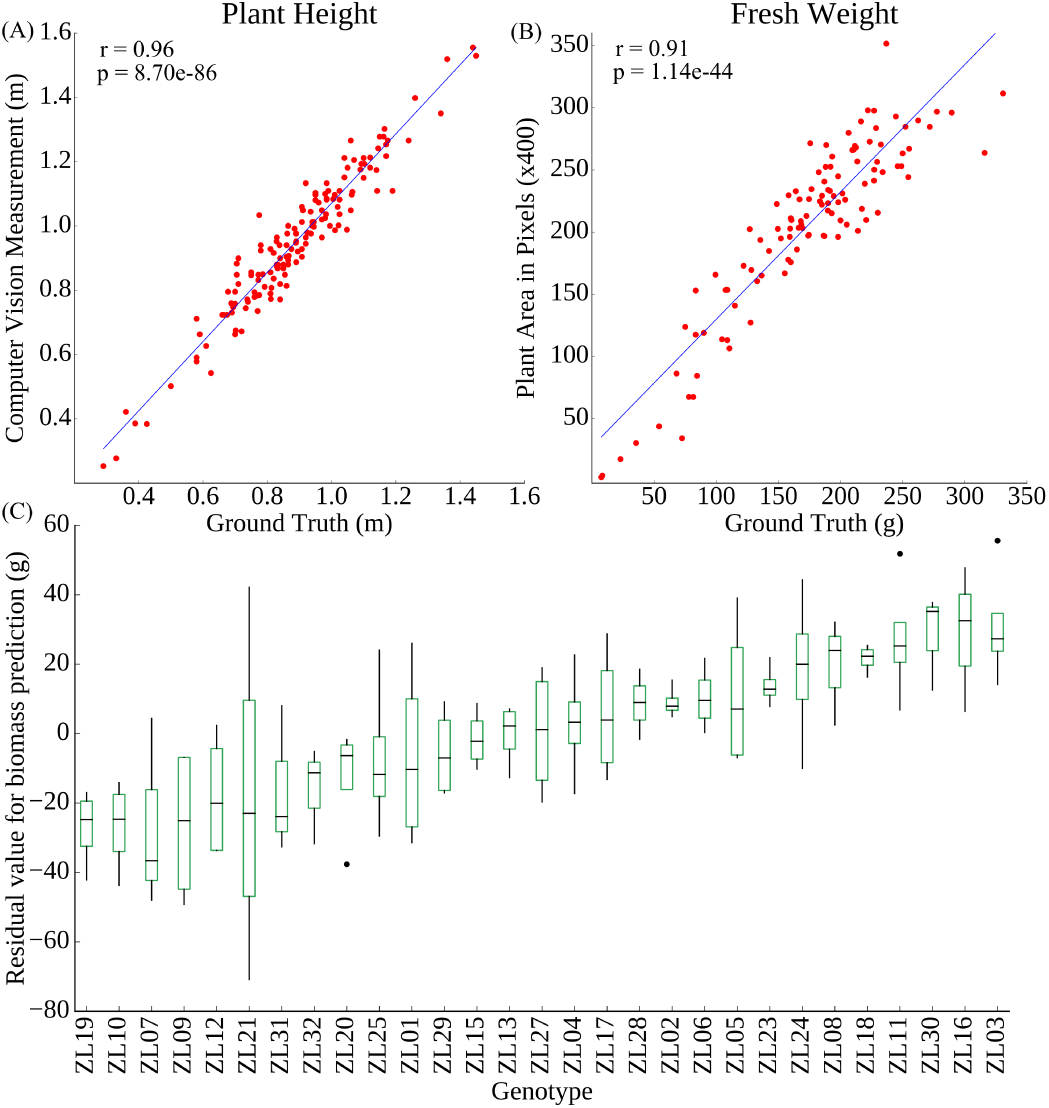
Correlation between image-based and manual measurements of individual plants.(A) Plant height; (B) Plant fresh biomass; (C) Variation in the residual between estimated biomass and ground truth measurement of biomass across inbreds.

The residual value – difference between the destructively measured biomass value and the predicted biomass value based on image data and the linear regression line equation – was calculated for each individual plant (Figure 2C). Using data from the multiple replicates of each individual accession, the proportion of error which is controlled by genetic factors rather than random error can be ascertained. We determined that 58% of the total error in biomass estimate was controlled by genetic variation between different maize lines. As such, this error is systematic rather than random and thus more likely to produce misleading downstream results when used in quantitative genetic analysis. As mentioned above, biomass and plant size are imperfectly correlated, as different plants can exhibit different densities, for example as a result of different leaf to stem ratios. Recent reports have suggested that estimates of biomass incorporating multiple traits extracted from image data can increase accuracy^14^. We tested the accuracy of biomass prediction of four multivariate estimation techniques on this dataset (see Methods). The correlation of the estimated biomass measures with ground truth data was 0.949, 0.958, 0.925 and 0.951 for multivariate linear model, MARS, Random Forest and SVM respectively. However, even when employing the most accurate of these four methods (MARS), 63% of the error in biomass estimation could be explained by genetic factors. This source of error, with the biomass of some lines systematically underestimated and the biomass of other lines systematically overestimated presents a significant challenge to downstream quantitative genetic analysis. Given the prevalence of plant pixel counts as a proxy for biomass^9,12,19–22^.

### Patterns of change over time

One of the desirable aspects of image based plant phenotyping is that, unlike destructively measured phenotypes, the same plant can be imaged repeatedly. Instead of providing a snapshot in time this allows researchers to quantify rates of change in phenotypic values over time, providing an additional set of derived trait values. Given the issues with biomass quantification presented above, measurements of plant height were selected to validate patterns of change in phenotypic values over time. As expected, height increases over time, and the patterns of increase tended to cluster together by genotype (Figure 3A). Increases in height followed by declines, as observed for ZL26, were determined to be caused by a change in the angle of the main stalk. While the accuracy of height estimates was assessed by comparison to physical ground truth measurements only on the last day, the height of three randomly selected plants (Plant 007-26, Plant 002-7 and Plant 041-29) were manually measured from image data and compared to software based height estimates, and no significant differences were observed between the manual and automated measurements (Figure 3B; Supplementary Table 1). To perform a similar test of the accuracy of biomass estimation at different stages in the maize life cycle, a set of existing ground truth measurements for two genotypes under two stress treatments^12^ were combined with additional later grow stage data (Supplemental Table 2). The correlation between total plant pixels observed in the two side views and plant biomass was actually substantially higher in this dataset (r=0.97) than the primary dataset, likely as a result of the smaller amount of genetic variability among these plants (Supplementary Figure 1).

**Figure 3.**
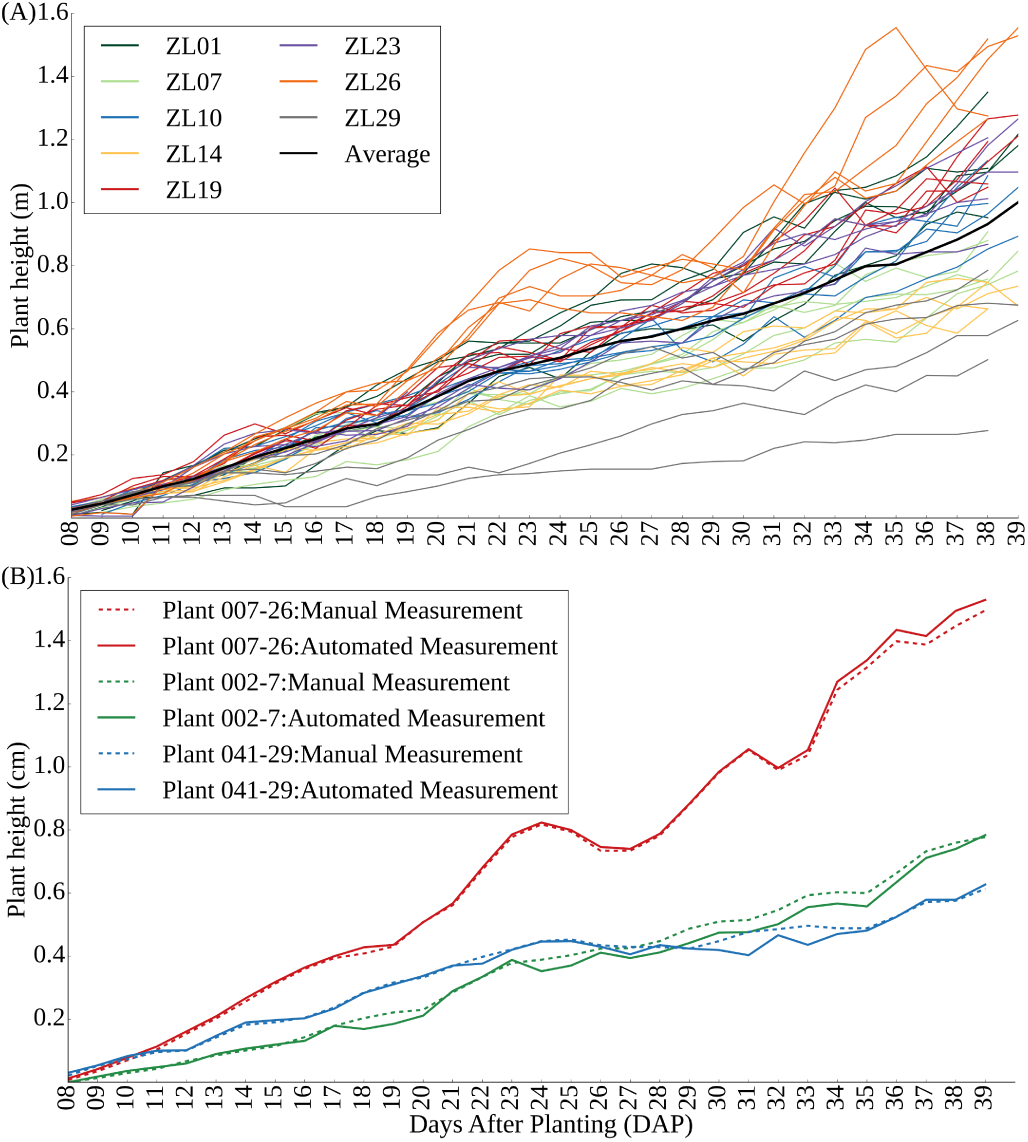
(A) Plant growth curves of each of five replicates of eight selected genotypes; (B) Comparison of manual measurements of plant height from image data with automated measurements for three randomly selected plants on each day of the experiment.

### Heritability of phenotypes

The proportion of total phenotypic variation for a trait controlled by genetic variation is referred to as the heritability of that trait and is a good indicator of how easy or difficult it will be to either identify the genes which control variation in a given trait, or to breed new crop varieties in which a given trait is significantly altered. Broad-sense heritability can be estimated without the need to first link specific genes to variation in specific traits^24^. Variation in a trait which is not controlled by genotype can result from environmental effects, interactions between genotype and environment, random variance, and measurement error. Controlling for estimated row effects on different phenotypic measurements significantly increased overall broad sense heritability (Figure 4A,B). This result suggests that even within controlled environments such as greenhouses, significant micro-environmental variation exists and that proper statistically based experimental design remains critical importance in even controlled environment phenotyping efforts.

**Figure 4.**
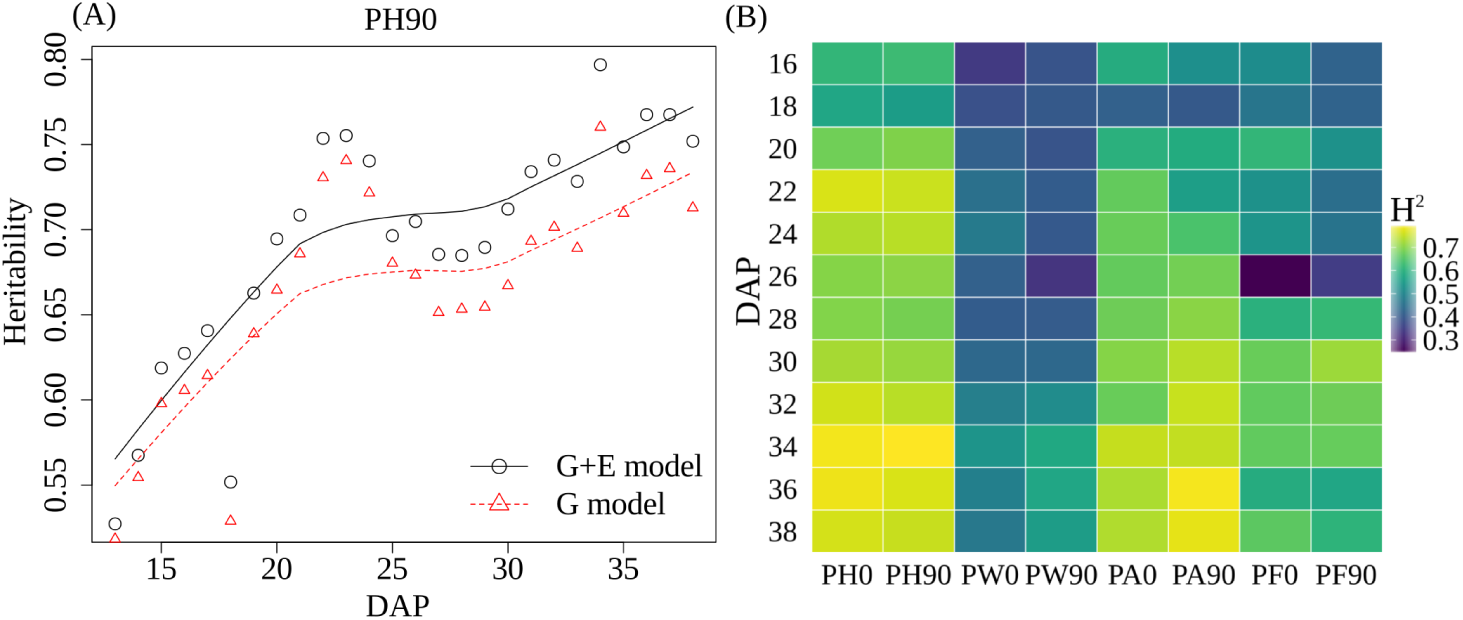
(A) The time course broad sense heritability of PH90. The heritability in the G model was calculated using a linear model that only considers the effect of genotype with residual values in the error term while heritability in the G + E model was calculated using a linear model that considers the effect of both genotype and environment (row effect) with residual values in the error term.; (B) The time course broad sense heritability of PA90 before and after controlling for the row effect; (B) Variation in broad-sense heritability (*H*^2^) after controlling row effects for 6 trait measurements every second day across the phenotyping cycle. PA0: Plant Area in 0 degree (The major axis of leaf phylotaxy was parallel to the camera at 0 degree); PA90: Plant Area in 90 degree (The major axis of leaf phylotaxy was perpendicular to the camera at 90 degree); PH0: Plant Height in 0 degree; PH90: Plant Height in 90 degree; PW0: Plant Width in 0 degree; PW90: Plant Width in 90 degree; PF0: Average of plant fluorescence intensity in 0 degree; PF90: Average of plant fluorescence intensity in 90 degree.

If the absolute size of measurement error was constant in this experiment, as the measured values for a given trait became larger, the total proportion of variation explained by the error term should decrease and, as a result, heritability should increase as observed (Figure 4A). This trend was indeed observed across six different phenotypic measurements (three traits calculated from each of two viewing angles (Figure 4B). Plant height also exhibited significantly greater heritability than plant area or plant width and greater heritability when calculated solely from the 90 degree side angle photo than when calculated solely from to 0 degree angle photo.

In previous studies, fluorescence intensity has been treated as an indicator for plant abiotic stress status^7,25–27^ or chlorophyll content level^28,29^. Using the fluorescence images collected as part of this experiment, the mean fluorescence intensity value for each plant image was calculated (see Methods). We found that this trait exibited moderate heritability, with the proportion of variation controlled by genetic factors increasing over time and reaching approximately 60% by the last day of the experiment (Figure 4B).

### Hyperspectral image validation

Hyperspectral imaging of crop plants has been employed previously in field settings using airborne cameras^30–32^. As a result ofthe architecture of grain crops such as maize, aerial imagery will largely capture leaf tissue during vegetative growth, and either tassels (maize) or seed heads (sorghum, millet, rice, oats, etc) during reproductive growth. The dataset described here includes hyperspectral imagery taken from the side of individual plants, enabling quantification of the reflectance properties of plant stems in addition to leaf tissue.

Many uses of hyperspectral data reduce the data from a whole plant or whole plot of genetically identical plants to a single aggregate measurement. While these approaches can increase the precision of intensity measurements for individual wavelengths, these approaches also sacrifice spatial resolution and can in some cases produce apparent changes in reflectivity between plants that result from variation in the ratios of the sizes of different organs with different reflective properties. To assess the extent of variation in the reflectance properties of individual plants, a principal component analysis of variation in intensity values for individual pixels was conducted. After non-plant pixels were removed from the hyperspectral data cube (Figure 5A) (See Methods), false color images were generated encoding the intensity values of the first three principal components of variation as the intensity of the red, green, and blue channels respectively (Figure 5B, C and D). The second principal component (green channel) marked boundary pixels where intensity values likely represent a mixture of reflectance data from the plant and from the background. The first principal component (red channel) appeared to indicate distinctions between pixels within the stem of the plant and pixels within the leaves.

**Figure 5.**
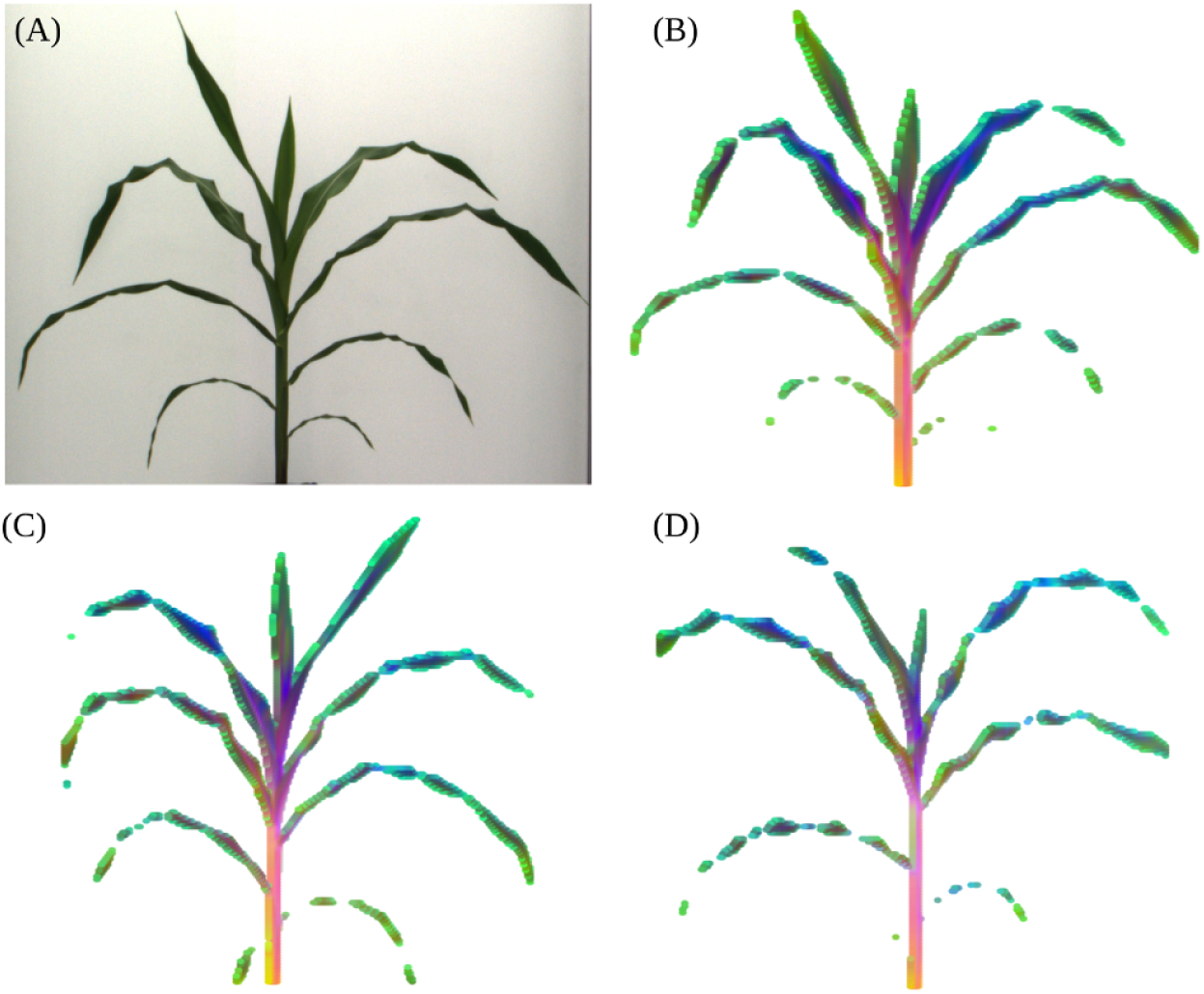
Segmentation and visualization of variation in hyperspectral signatures of representative maize plant images. (A) RGB photo of Plant 013-2 (ZL02) collected on DAP 37; (B) False color image constructed of the same corn plant from a hyperspectral photo taken on the same day. For each plant pixel the values for each of the first three principal components of variation across 243 specific wavelength intensity values are encoded as one of the three color channels in the false image; (C) Equivalent visualization for Plant 048-9 (ZL09); (D) Equivalent visualization for Plant 008-19 (ZL19).

Based on this observation, an index was defined which accurately separated plant pixels into leaf and stem (see Methods). Stem pixels were segmented from the rest of the plant using an index value derived from the difference in intensity values observed in the 1056nm and 1151nm hyperspectral bands. This methodology was previously described^12^. The reflectance pattern of individual plant stems is quite dissimilar from the data observed from leaves and exhibits significantly different reflective properties in some areas of the near infrared (Figure 6). Characteristics of the stem are important breeding targets for both agronomic traits (lodging resistance, yield for biomass crops) and value added traits (biofuel conversion potential for bioenergy crops, yield for sugarcane and sweet sorghum). Hyperspectral imaging of the stem has the potential to provide nondestructive measurements of these traits. The calculated pattern of leaf reflectance for the data presented here are comparable with those observed in field-based hyperspectral studies^33–35^, providing both external validation and suggesting that the data presented here may be of use in developing new indices for use under field conditions.

**Figure 6.**
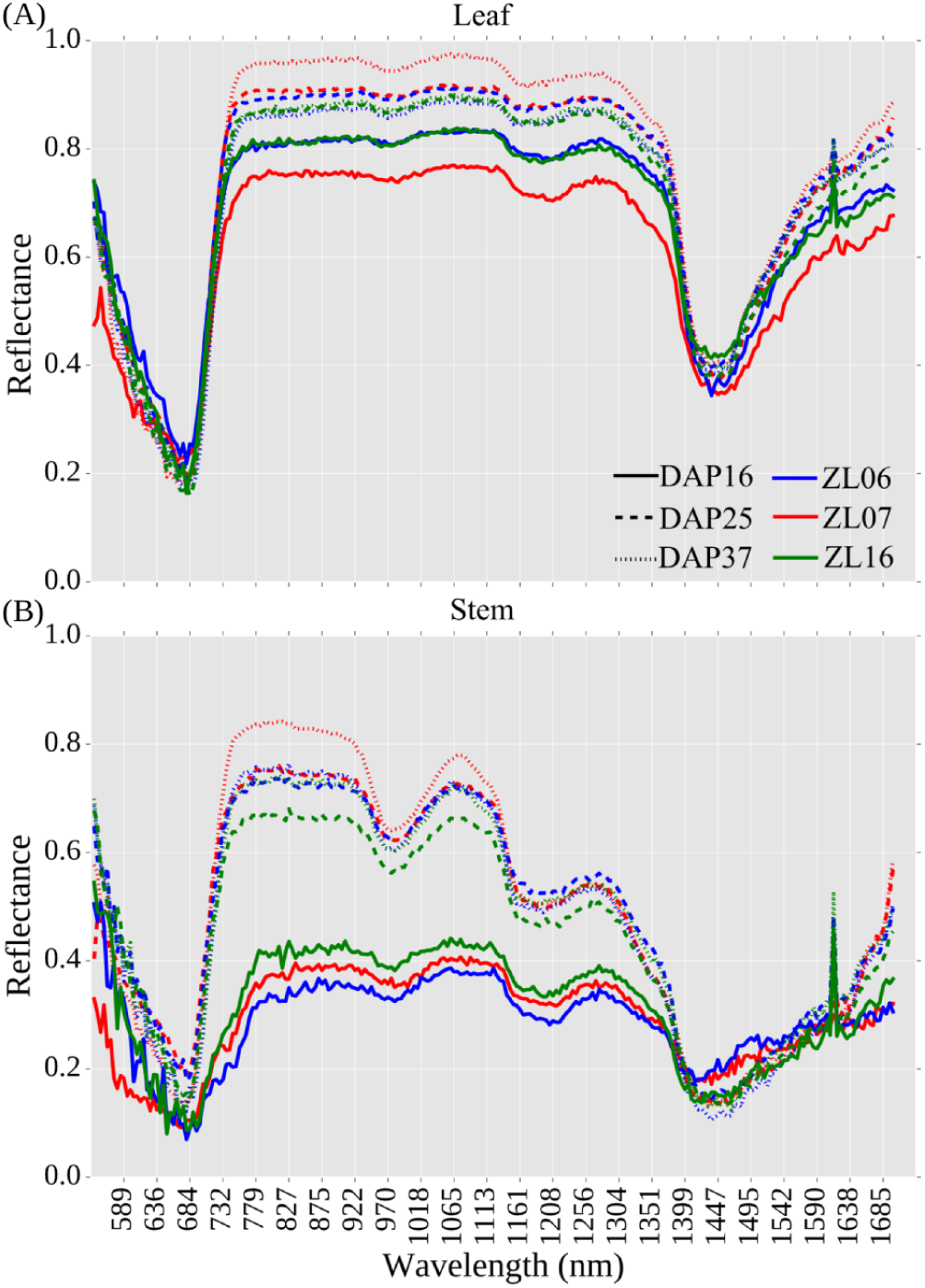
Reflectance values for three plants - Plant 090-6 (ZL06), Plant 002-7 (ZL07), and Plant 145-16 (ZL16) on three days across development. (A) Reflectance values for non-stem plant pixels (i.e. leaves) (B) Reflectance values for pixels within the plant stem.

In conclusion, while the results presented above highlight some of the simplest traits which can be extracted from plant image data, these represent a small fraction of the total set of phenotypes for which image analysis algorithms currently exist, and those in turn represent a small fraction of the total set of phenotypes which can potentially be scored from image data. Software packages already exist to measure a range of plant architectural traits such as leaf length, angle, and curvature from RGB images^6,36^. Tools are also being developed to extract phenotypic information on abiotic stress response patterns from fluorescence imaging^6,7^. The analysis of plant traits from hyperspectral image data, while common place in the remote sensing realm where an entire field may represent a single data point, is just beginning for single plant imaging. Recent work as highlighted the potential of hyperspectral imaging to quantify changes in plant composition and nutrient content throughout development^12,37^. While these techniques have great potential to accelerate efforts to link genotype to phenotype through ameliorating the current bottleneck of plant phenotypic data collection, it will be important to balance the development of new image analysis tools with the awareness of the potential for systematic error resulting from genetic variation between different lines of the same crop species.

**1 Availability of source code and requirements**

- Project name: Maize Phenotype Map
- Project home page: https://github.com/shanwai1234/Maize_Phenotype_Map
- Operating system(s): Linux
- Programming language: Python 2.7
- Other requirements: OpenCV module 2.4.8, Numpy *>*1.5, CMake *>* 2.6, GCC *>* 4.4.x, Scipy 0.13
- License: BSD 3-Clause License

## 2 Availability of supporting data and materials

The image data sets, pot weight records per day and ground truth measurements with corresponding documentations from 4 types of cameras of 32 maize inbreds and same types of image data for the two maize genotypes under two stress treatments were deposited in the CyVerse data commons under a CC0 license with doi: 10.7946/P22K7V. (The data for the peer review process can be downloaded from https://doi.org/10.7946/P22K7V). All image data were stored in the following data structure: Genotype *- >* Plant *- >* Camera type *- >* Day. For the hyperspectral camera each photo is stored as 243 sub images, each image representing intensity values for a given wavelength, so these require one additional level of nesting in the data structure Day *- >* wavelength. The grayscale images from the IR camera and the hyperspectral imaging system are stored as three-channel images with all three channels in a given pixel set to identical values. The fluorescence images contain almost all information in the red channel with the blue and green channel having intensities equal to or very close to zero, but data all three channels exist. Genotype data of 32 inbreds were generated as part of a separate project and can be retrieved from either https://doi.org/10.7946/P2V888 or http://cbsusrv04.tc.cornell.edu/users/panzea/download.aspx?filegroupid=4. Measurements for thirteen core phenotypes at each field trial as well as local weather data can be retrieved from publicly released Genomes 2 Fields datasets released on CyVerse. Data from the 2014 G2F field trials is posted (https://doi.org/10.7946/P2V888) and data from the 2015 G2F field trials is posted (https://doi.org/10.7946/P24S31). Genetically identical seeds from the majority of the accessions used in creating both this dataset and the genomes to fields field trial data can be ordered from public domain sources (e.g. USDA GRIN) and are listed in Table 1.

## 3 Declarations

### 3.1 List of abbreviations

DAP: Days after planting

GBS: Genotyping by Sequencing

LED: Light-emitting diode

MARS: Multivariate Adaptive Regression Splines

NDVI: Normalized difference vegetation index

NIR: Near-infrared

RGB: An image with separate intensity values for the red, blue and green channels

SNP: Single Nucleotide Polymorphism

SVM: Support Vector Machines

UNL: University of Nebraska-Lincoln

PA0: Plant Area calculated from a 0 degree image. Plants were initially orientated then leaves would be arranged parallel to the

camera at 0 degrees.

PA90: Plant Area calculated from a 90 degree image. Plants were initially orientated then leaves would be arranged perpendicular

to the camera at 90 degrees.

PCA: Principal Component Analysis

PH0: Plant Height calculated from a 0 degree image

PH90: Plant Height calculated from a 90 degree image

PW0: Plant Width calculated from a 0 degree image

PW90: Plant Width calculated from a 90 degree image

PF0: Average of plant fluorescence intensity in 0 degree

PF90: Average of plant fluorescence intensity in 90 degree.

### 3.2 Consent for publication

Not applicable.

### 3.3 Competing Interests

The authors declare that they have no competing interests.

### 3.4 Funding

This research was supported by the Nebraska Corn Board (Award #88-R-1617-03), the Iowa Corn Board (Award #), the National Science Foundation under Grant No. OIA-1557417, and Internal University of Nebraska funding to JCS. The sources of funding have no role in the design of the study and collection, analysis, and interpretation of data and in writing the manuscript.

### 3.5 Author’s Contributions

JCS, and YQ designed the experiment; VS, JCS and ZL performed data acquisition; ZL, PP, YQ, YX, YG and JCS analyzed and interpreted the data; ZL and JCS produced and curated the metadata; ZL and JCS implemented software; ZL and JCS prepared the initial draft. All authors reviewed the manuscript.

## 4 Acknowledgements

The authors are grateful to Yang Zhang, Xianjun Lai and Daniel WC Ngu for help in collecting manual measurements of plants, Thomas Hoban for manually counting pixels of selected plant images, Kent M. Eskridge for valuable discussions on experimental design, Addie Thompson, Jinliang Yang for assistance on heritability analysis, and the members of the Genomes to Fields consortium for sharing both seed and datasets prior to publication. CyVerse is supported by the U.S. National Science Foundation under award numbers DBI-0735191 and DBI-1265383.

